# End-to-end genomic prediction: direct prediction of images and text from genome-wide molecular markers

**DOI:** 10.1101/2025.11.03.686395

**Authors:** Mark T. Watson, Mitchell Feldmann, Haipeng Yu, Hao Cheng

## Abstract

**Background:** Methods to predict the heritable component of phenotypes from genetic markers, collectively known as genomic prediction, have been widely applied in plant and animal breeding and in human genetics. Currently, genomic prediction is limited to numeric phenotypes. In some cases, though, plant and animal phenotypes are better understood through images and text rather than numbers. The current best practice for incorporating images and text in genomic prediction is to first extract scalar numeric phenotypes from images and text, and then to perform genomic prediction on the numeric phenotypes. While this approach is effective for some traits, it involves discarding most of the information in the image or text, including potentially useful information. Additionally, numeric phenotypes derived from images and text may not be as interpretable as the images and text themselves.

**Approach:** We present a novel approach for predicting images and text from SNP markers, which we refer to as end-to-end genomic prediction, and validate this approach using genotypes, text, and image phenotypes derived from a strawberry (*Fragaria × ananassa*) diversity panel. Our approach combines nonlinear latent space encoding with linear genomic prediction to generate accurate breeding values for a high-dimensional phenotype, for example, images or text.

**Results:** For both genome-to-image and genome-to-text prediction, we found that predicting images and text and then extracting numeric traits from them was in some cases as accurate as directly predicting extracted numeric phenotypes, demonstrating for the first time that genome-to-image prediction accuracy can be comparable to conventional genomic prediction accuracy.

**Conclusions:** Based on our proof-of-concept using the same core end-to-end method in both images and text, we believe end-to-end genomic prediction could be of use in a wide range of visual and multidimensional phenotypes in plants and animals, although further work to improve the accuracy of the embedding and genomic prediction steps is needed.

## Introduction

Genetic improvement of agricultural populations is vital for enhancing the efficiency, sustainability, and profitability of breeding programs. By selecting plants or animals with desirable traits, such as higher yields or disease resistance, as parents, breeders can achieve permanent and cumulative genetic improvements within their populations. The success of genetic improvement hinges on a core question: how accurately can we predict breeding values (the additive genetic value of an individual) to identify and select the most promising candidates for selective breeding?

The discovery of genome-wide high-density molecular markers (e.g., single-nucleotide polymorphisms, SNPs) and effective ways to assay them has revolutionized genetic improvement of complex traits in agriculture^[9,17,19,37,38,42]^. This progress is mainly driven by genomic prediction^[23]^, a statistical method using genome-wide molecular markers to predict breeding values. Genomic prediction can be framed as a supervised learning problem. In this context, the labeled dataset consists of input-output pairs, where the input represents the genome-wide molecular markers, and its associated output (label) is the phenotype.

Plant and animal breeding programs invest significant resources in collecting non-numeric phenotypes, such as images and text, as part of trait assessment for selection and for recordkeeping apart from selection^[12,21,24,32]^. Technological advancements over recent decades have expanded the variety and complexity of data types collected, enabling breeders to capture a broader range of phenotypic traits. A wide range of light-based sensors, from smartphone cameras to aerial hyperspectral cameras, are now employed to capture detailed images in plant and animal populations^[1,4,10,16,31,33,45]^. These images are typically then converted into numeric values such as subjective ratings and shape index for analysis. Text data, although less commonly used, plays a critical role in documenting non-standardized and subjective traits, particularly for aesthetic, sensory, and behavior evaluations^[6,11,27,46]^. The limited use of text data stems from the challenge of robustly translating nuanced descriptions into quantifiable metrics, and then treating those as numeric phenotypes for conventional genomic prediction.

Currently, non-numeric phenotypes like images are usually converted to numeric phenotypes before genomic prediction analysis^[20,43]^. However, without appropriate data analysis methods, these approaches can lead to data being underutilized or misinterpreted. Extracted numeric traits from images or text generally don’t account for the full variation present in the higher-dimensional phenotype, and human bias is introduced in the choice of which features to extract^[15]^. Given the unique properties of image and text phenotypes, suitable prediction methods need to be developed for these phenotypes to enable their full use in breeding programs. Analogously, it was the combination of data from genome-wide molecular markers with suitable analytical methods like mixed effect models that enabled the advancement of genomic prediction. Analytical methods must be carefully adapted to address the specific challenges of analyzing non-numeric data, such as images and text, to ensure the data’s full potential is realized in breeding applications.

So far, genomic prediction has been restricted to scalar numeric phenotypes, influencing the way traits are analyzed and, at times, limiting interpretability. While numerical values are suitable for certain traits, such as yield or body weight, complex phenotypes are often better represented through images or text. Traits like shape, color, and disease symptoms are inherently visual, and reducing these to numerical scores can strip away critical information, potentially compromising the accuracy of selection decisions^[15]^. A recent method called GenoDrawing has been proposed to predict images solely from genetic markers^[22]^. However, this approach is limited by its inability to effectively use genome-wide genetic markers and lacked a direct comparison with conventional genomic prediction methods.

Here we introduce End-to-End Genomic Prediction, a framework for directly predicting both images and text from genome-wide genetic markers. Specifically, we use linear mixed models to predict numeric representations of images that are sufficient to accurately reconstruct images. This means our prediction target remains the original endpoint (images or text) unlike in existing approaches where the endpoint is changed to simplified scalar phenotypes to facilitate prediction. Latent space encodings have been used to simplify complex traits in strawberries and other species, but have not previously been combined with linear mixed models for genomic prediction^[22,30,35,47]^. End-to-end genomic prediction circumvents the current requirement of converting images and text into numeric values, a step that often diminishes interpretability and reduces the information density inherent in their original forms. This limitation has historically hindered the incorporation of images and textual data into genomic prediction studies. Our end-to-end approach synergistically integrates flexible machine learning techniques, such as neural network-based latent space encoding, with classical statistical methods like linear mixed models, enhancing both the interpretability and practical utility of genomic predictions.

## Methods

### End-to-end genomic prediction

The same core end-to-end genomic prediction method underlies both our genome-to-image and genome-to-text predictions, combining neural network-based latent space encoding with conventional genomic prediction using linear mixed models. A schematic of our method is shown in Figure 1. In the latent space encoding step, we use a neural network capable of converting high-dimensional image or text phenotypes into embeddings in a potentially lower-dimensional continuous latent space, from which the original data can be accurately reconstructed. Then in the genome-to-embedding prediction step, we trained a linear mixed model to predict these embeddings from genotype data. Conceptually, this approach parallels the Neural Network Mixed Model (NNMM)^[44]^, which integrates neural networks with mixed models to incorporate intermediate, biologically meaningful features within the genomic prediction framework. In our case, these intermediate features correspond to image and text embeddings, serving as structured representations that bridge complex phenotypes and genomic variation. After training, in the full end-to-end genomic prediction method, we predict embeddings directly from genotypes and decode them back into images or text using the decoder component of the variational autoencoder. Thus, in the end-to-end method, breeding values for images or text can be predicted for individuals for which these high-dimensional traits have not been directly observed.

**Figure 1.**
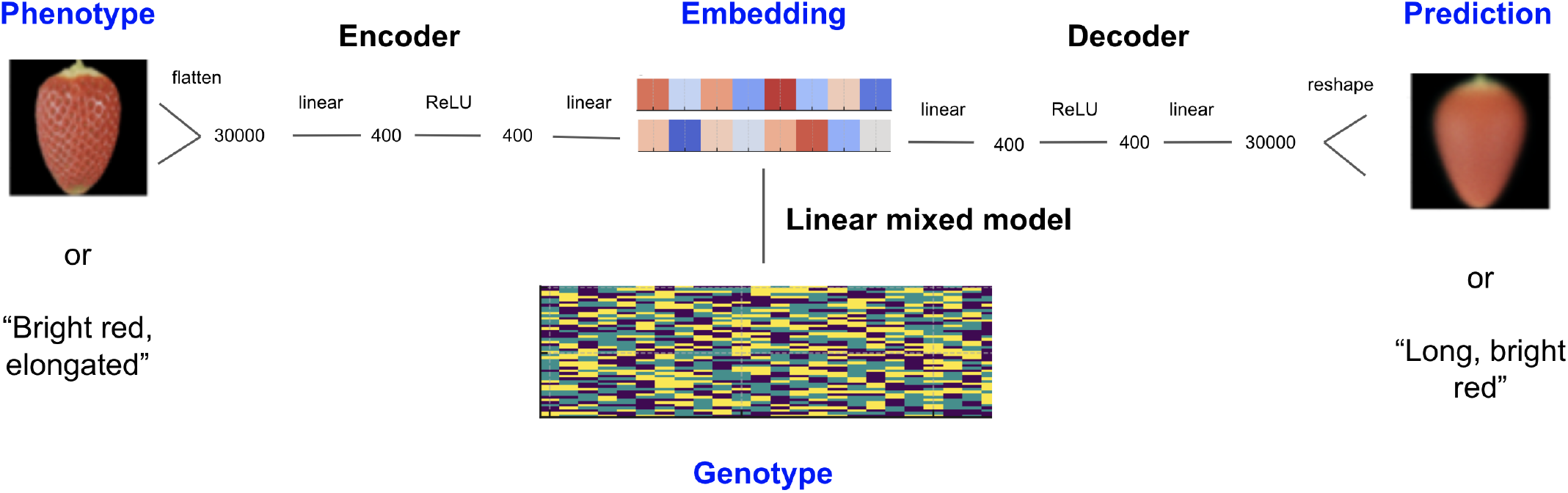
Schematic of our end-to-end genomic prediction method. This method consists of two main components: nonlinear latent space encoding and genome-to-embedding prediction. In the latent space encoding step, a neural network is trained to reduce high-dimensional phenotypes to latent space embeddings with high reconstruction accuracy. Then, in the genome-to-embedding prediction step, a linear mixed model is trained to predict embeddings of a high-dimensional phenotype (for example, images or text). The latent space encoding step is then used again in reverse to decode predicted embeddings to images or text. The neural network architecture and matrix / vector dimensions shown are for genome-to-image prediction.

### Description of image, text, and genotype data

For both our genome-to-image and genome-to-text prediction analysis, we used a strawberry fruit image and genotype dataset previously collected by the Strawberry Breeding and Research Program at UC Davis, paired with text descriptions we derived from the image dataset. The dataset represents a population of 563 strawberry parental and hybrid clones with a broad range of phenotypic diversity for heritable fruit traits, as described in^[14]^. Strawberry clones were grown in Salinas, CA, in a randomized complete block design with three replicates of each genotype, and they were harvested over two years.

Harvested fruit were imaged with the calyx removed, following the protocol described in^[14]^. We segmented individual berries from raw images using OpenCV^[5]^ code in Python adapted from^[41]^, ensuring that images had a consistent size (100 pixels x 100 pixels x 3 RGB channels) and rotation (calyx side facing up). After filtering out berries that were improperly segmented, the dataset contained 13,922 segmented strawberry images, with each clone represented by an average of 24 images. We derived text phenotypes from the image dataset using the method described in the genome-to-text prediction methods section. The resulting text ratings adhered to the prompt and used similar vocabulary as subjective human ratings while reflecting variation in shape and color from the image dataset. All clones were genotyped for SNPs using the Axiom FanaSNP Strawberry 50K Genotyping Array^[18]^.

### Genome-to-image prediction

For genome-to-image, we performed an experiment to test whether genome-to-image prediction accuracy would be comparable to the prediction accuracy of conventional genomic prediction. We used scalar numeric traits as a means of comparing end-to-end genomic prediction with conventional genomic prediction. These traits, referred to as “extracted traits,” are derived from images using image analysis, following current best practices for incorporating images in genomic prediction. The five extracted traits we selected, which have been used in strawberry and other species^[3,8,14,36,47]^, are defined below:

1. Length-width ratio (LWR): the ratio of the longer dimension of a berry to its shorter dimension.
2. R: the mean red color component of a berry in RGB space^[34]^.
3. G: the mean green color component of a berry in RGB space.
4. B: the mean blue color component of a berry in RGB space.
5. Redness: the Euclidean distance, in CIELAB color space^[28]^, between the average color of a berry and an ideal red berry color.

We used CIELAB space for the redness metric due to its improved perceptual uniformity over RGB space. Each extracted trait was calculated as the mean across all strawberry images for each genotype to account for the substantial variability between berries within a genotype. We created an 80:20 train-test split by genotype, ensuring that multiple images from the same genotype were consistently placed within either the training or testing set. We generated fifty distinct train-test splits using Monte Carlo resampling, and then compared end-to-end and conventional genomic prediction on each train-test split.

For each train-test split, we trained a variational autoencoder (VAE) implemented in Pytorch^[29]^ on the images in the training set. We obtained model accuracy for the VAE step by itself by comparing the extracted traits of the original images to those of the encoded and subsequently decoded images. We used a simple VAE architecture with an embedding size of 16 we used for image embeddings, as shown in 1. Hyperparameters including embedding size, learning rate, and batch size were optimized using grid search prior the main experiment, which used the optimal hyperparameters we identified. We then regressed the embeddings generated by the VAE (dependent variables) on genotypes (independent variables; aa, Aa, AA = 0, 1, 2 for biallelic SNPs under an additive genetic model) using ridge-regression best linear unbiased prediction (RR-BLUP), implemented via the R package rrBLUP^[13]^.

Using the same 80:20 train-test splits as before, we predicted embeddings from genotypes, which we then used to generate predicted images. To evaluate end-to-end prediction accuracy, we compared the extracted traits from these predicted images to the ground truth extracted traits (see Figure 2). To be able to compare end-to-end genomic prediction accuracy with conventional genomic prediction accuracy, we assessed the accuracy of conventional genomic prediction by directly predicting the extracted traits without the intermediate image step.

**Figure 2.**
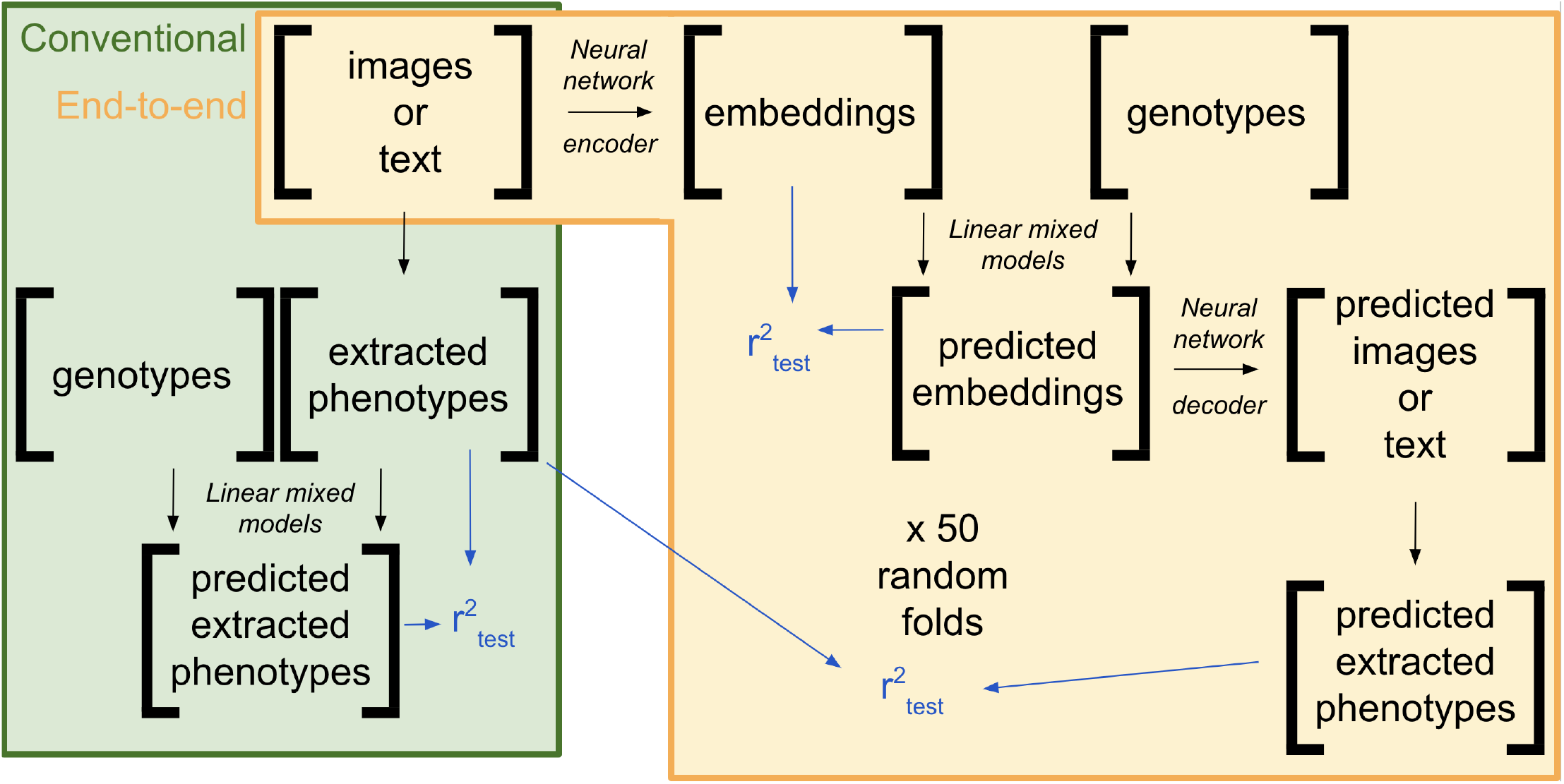
Validation scheme used to compare genome-to-image prediction accuracy to conventional genomic prediction. In this scheme, we use scalar, biologically relevant “extracted phenotypes” such as length-width ratio and redness as a consistent metric to compare the two methods, since conventional genomic prediction cannot effectively predict images. A training set is used to train autoencoder and genome-to-embedding prediction models, which are then used to end-to-end genomic prediction in a testing set. Since the test set prediction accuracy for extracted phenotypes for conventional and end-to-end genomic prediction represent the same overall prediction task (genome-to-extracted-traits), we compare them as the best means for evaluating the prediction task we are most interested in but unable to compare across methods (genome-to-image). We used the same validation scheme for our genome-to-text experiment, substituting text phenotypes and text embeddings for image phenotypes and image embeddings, and using a pretrained embedding model (text-embedding-ada-002).

### Genome-to-text prediction

For the genome-to-text experiment, due to the lack of human-derived text phenotypes in the strawberry dataset, we generated text phenotypes from the five image-derived traits (LWR, R, G, B, and redness) by prompting a large language model (OpenAI’s GPT-4o-mini) with a string of the five image-derived traits and the following prompt: “Based on the provided phenotype data, describe the strawberry in 3-4 words. For reference, an LWR of 1 is short and LWR of 1.5 is long, and a redness of 80 is pale, a redness of 130 is light red, and a redness of 180 is deep red. R ranges from 30 to 100, G ranges from 50 to 140, B ranges from 140 to 240, and all three are inversely correlated with Redness.” Based on the generated text phenotypes, we compared genome-to-text prediction with conventional genomic prediction. The core end-to-end framework was the same as that used for genome-to-image prediction. However, instead of training an autoencoder to convert text to embeddings, we used OpenAI’s pre-trained text-embedding-ada-002 model to generate vector embeddings for all text phenotypes. These embeddings were then predicted from genotype data using rrBLUP. To decode the predicted embeddings back into text, we used the vec2text embedding inversion model^[25,26]^, trained specifically for textembedding-ada-002.

To compare genome-to-text with conventional genomic prediction, we converted the text phenotypes to numeric traits. Specifically, we extracted two traits, including length and redness, due to the fact that text phenotypes did not contain enough information to reconstruct exact RGB color components. These traits were extracted by prompting GPT-4o-mini with the following instruction: “Based on the provided phenotype description, generate numeric values for length and redness. Length should range from 0 (short) to 1 (long), and redness should range from 0 (not red) to 1 (red). Your response should be strictly of the form ‘Length:, Redness:’”. To estimate end-to-end prediction accuracy, we compared extracted traits from these predicted images to the ground truth extracted traits (see Figure 2).

## Results

### Genome-to-image prediction

#### Evaluating the performance of latent space encoding and genome-to-embeddings prediction

Table 1 shows the reconstruction for strawberry fruit images using VAE embeddings, based on encoding images using the autoencoder and then decoding the resulting embeddings back to images. We identified an optimal VAE embedding size of 16 by testing various sizes from 1 to 256, observing that reconstruction accuracy plateaued around an embedding size of 16 for our dataset. Reconstruction accuracy is measured as mean squared error (MSE) for pixel features and squared Pearson correlation *r*^2^ between true and predicted values for 5 extracted traits (LWR, R, G, B, and redness). Reconstruction accuracy was nonzero and moderately high (*r*^2^ > 0.5) across all 50 replicates for pixel features and all 3 extracted traits. This indicates that the variational autoencoder preserved the general shape and color of the image. However, as prediction accuracy was also less than 1 for all extracted traits, some relevant information was lost during latent space encoding using our current approach. Some of this loss in information may be attributable to blurring through local averaging in the variational autoencoder, which has previously been described as a characteristic of VAEs ^[7]^. A comparison between original and VAE decoded mean embedding images is included in Figure 4.

**Table 1.** Test set pixel level mean squared error (MSE) and prediction accuracy (*r*^2^) for extracted traits as measures of autoencoder reconstruction accuracy. Prediction accuracy was moderately high (*r*^2^ > 0.5), which indicates that the autoencoder was able to encode breeding-relevant shape and color.

| Trait | MSE (sd) | $r^2$ (sd) |
| --- | --- | --- |
| Pixel | 68.3 (0.392) | – |
| LWR | – | 0.790 (0.012) |
| R | – | 0.752 (0.017) |
| G | – | 0.850 (0.013) |
| B | – | 0.673 (0.025) |
| Redness | – | 0.855 (0.011) |

In the genome-to-embedding prediction step, the VAE embedding components had moderate heritability and prediction accuracy in replicates, with a mean test set prediction accuracy across embedding components of 0.247 and a standard deviation of 0.03. Genome-to-embedding prediction accuracy was higher than genomic prediction accuracy for a random sample of pixels that varied between images, which had an average of 0.220. This suggests that image embedding components are a more appropriate prediction target for genome-to-image prediction than pixels. Additionally, genomic prediction of raw pixels did not produce meaningful image outputs in spite of the nonzero prediction accuracy due to extensive blurring in the predicted image beyond the level of blurring in autoencoder-based prediction.

#### Comparing genome-to-image prediction and conventional approaches

Table 2 presents the prediction accuracies for the extracted traits using our end-to-end genome-to-image method and conventional genomic prediction. For R, G and Redness, end-to-end prediction accuracy was similar to conventional genomic prediction accuracy, demonstrating for the first time that genome-to-image prediction can match or even exceed conventional genomic prediction accuracy. For LWR and B, the endto-end prediction accuracy was lower than the conventional genomic prediction accuracy.

**Table 2.** Comparison between accuracy (r^2^) for end-to-end genome-to-image prediction and conventional genomic prediction for extracted traits.

|  | Prediction accuracy (sd) |  |
| --- | --- | --- |
|  | end-to-end | conventional |
| LWR | 0.074 (0.048) | 0.159 (0.054) |
| R | 0.257 (0.064) | 0.330 (0.056) |
| G | 0.316 (0.085) | 0.383 (0.077) |
| B | 0.073 (0.048) | 0.281 (0.065) |
| Redness | 0.339 (0.072) | 0.377 (0.077) |

Figure 3 shows the difference between end-to-end prediction accuracy (r^2^) and conventional genomic prediction accuracy (r^2^) for genome-to-image prediction for five extracted traits. Each point within a distribution represents the difference calculated for a single experimental run (out of 50) on one trait. A difference of less than 0 indicates that end-to-end genomic prediction performed worse than conventional genomic prediction for that trait on that run, while difference greater than 0 indicates that end-to-end genomic prediction performed better than conventional genomic prediction for that trait on that run. End-to-end prediction accuracy was substantially higher than 0 and slightly lower than conventional genomic prediction accuracy for all traits, but not significantly for a difference of 0.1%. Additionally, end-to-end prediction accuracy was higher than end-to-end prediction accuracy for some replicates.

**Figure 3.**
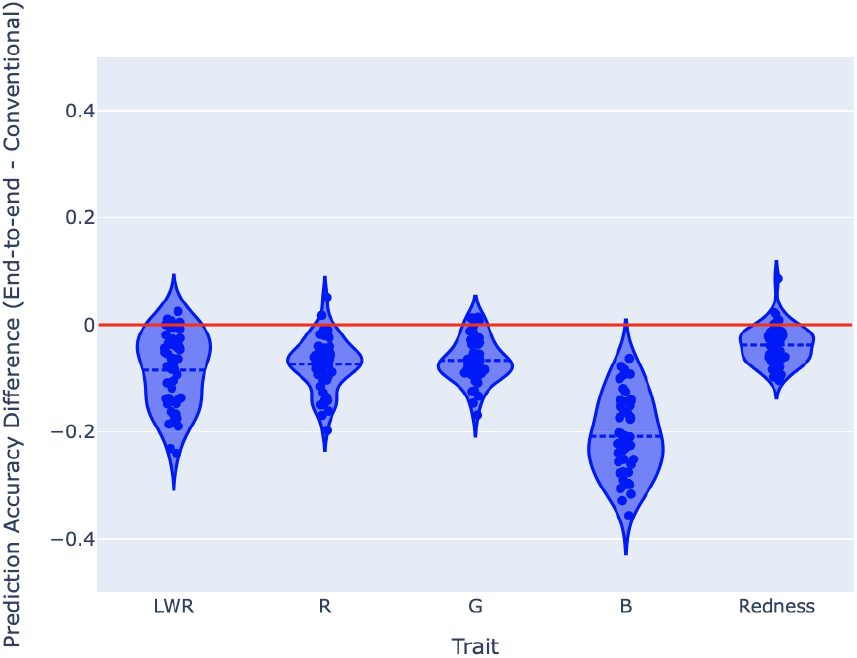
Violin plot of the difference between end-to-end prediction accuracy (r^2^) and conventional genomic prediction accuracy (r^2^) for genome-to-image prediction, as measured by five extracted traits.

#### Visualizing the genome-to-image prediction pipeline across genotypes

Figure 4 compares the outputs from each step of our genome-to-image method. Each column represents a different stage in our end-to-end method applied to genome-to-image prediction: original images, decoded mean embedding images, mean decoded mean embedding image, and end-to-end predicted image. Each row represents one clone. To demonstrate our end-to-end method’s ability to predict shape and color traits accurately, we picked four clones from the testing set of the first replicate which demonstrate widely varying combinations of shape and color: the first clone is generally long and tapered with red color, the second clone is short and round with an orange-red color, the third clone is wide and tapered with a deep red color, and the fourth clone is deep red and with less taper. The first column shows the range of image phenotypes observed for these clones; while variation exists within each clone, there is sufficient between-clone variation to consider some clones better than others for selection based on the images. For example, the light color and short shape of the second clone is less desirable than the dark color and tapered shape of the third clone. The second column shows the results of encoding each image and decoding it using a trained autoencoder. The general shape and color of each image is visibly well preserved, which is consistent with the high reconstruction accuracy for extracted shape and color traits shown in Table 1, although small scale features are lost through blurring. Since each clone is represented by multiple images, we took the mean of the embeddings for each clone prior to the embedding prediction step. The third column shows the decoded version of the mean embedding for each clone. In our experiment, we predicted an embedding for each clone from SNP markers. The fourth column shows the decoded results of the embedding predicted model, which are the end result of our end-to-end prediction method. The end-to-end predicted images are subjectively similar to the mean-decoded images, indicating that our end-to-end genomic prediction method predicts image phenotypes that are subjectively similar to the true image phenotypes. For example, the first clone is darker and more tapered than the second clone according to the mean decoded mean embedding images, and the end-to-end predicted image for the first clone is also darker and more tapered than the predicted image for the second clone. The end-to-end predicted image for the first clone is also much more similar to the mean decoded mean embedding image for the first clone than to the mean decoded mean embedding image for the second clone. Additional differences in color and tapering are also successfully predicted for the third and fourth clones. This correspondence between our end-to-end prediction target and results is discussed above and shown quantitatively in Table 2.

**Figure 4.**
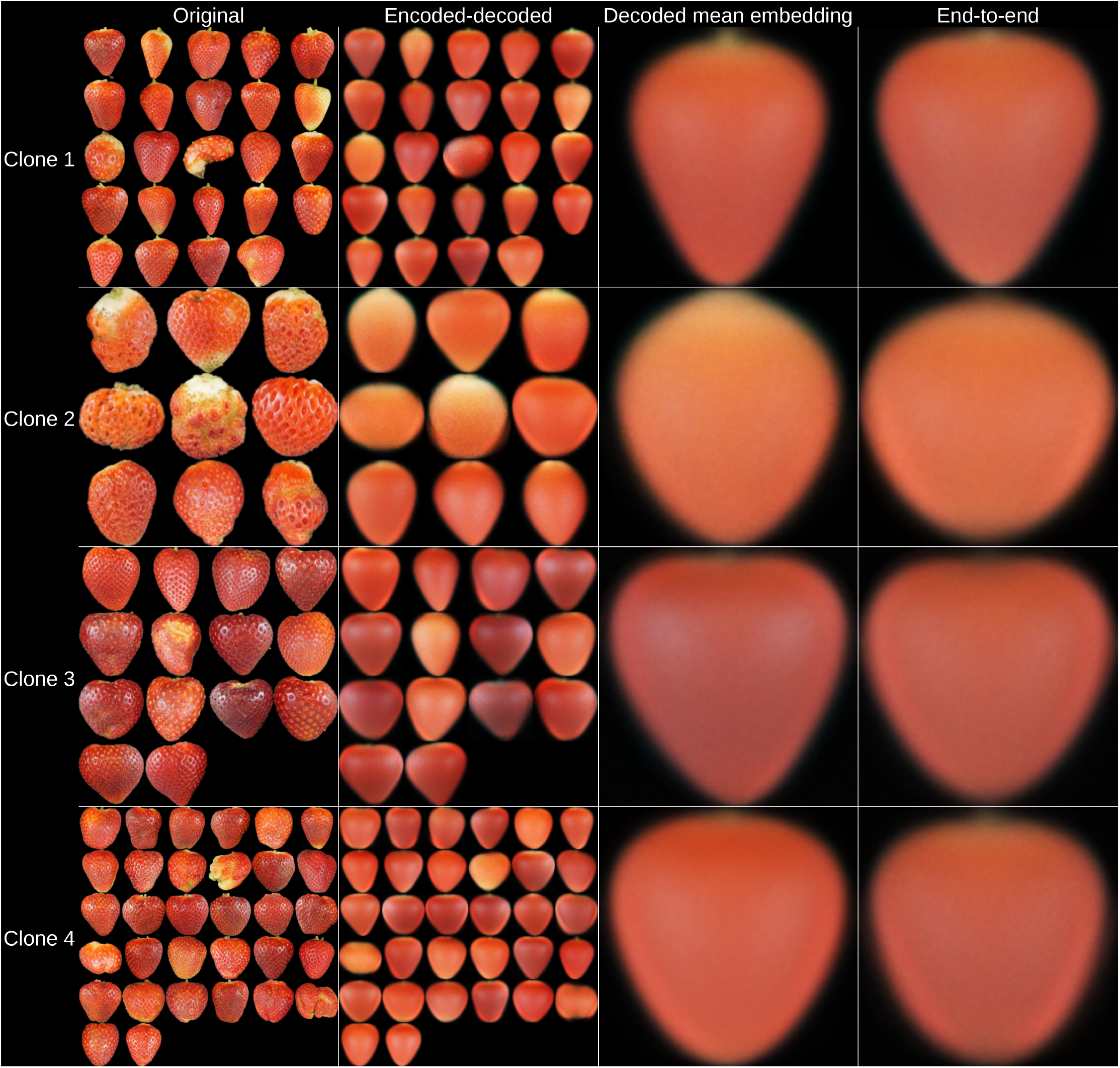
A comparison between genome-to-image outputs. The four columns represent original images, decoded mean embedding images, mean decoded mean embedding image, and end-to-end predicted image for four test set clones picked to represent the most divergent combinations of shape and color in the population. Each row represents one strawberry clone.

### Genome-to-text prediction

#### Evaluating the performance of latent space encoding and genome-to-embeddings prediction

The pretrained text embedding model we used (text-embedding-ada-002 for encoding paired with vec2text for decoding) had very high reconstruction accuracy, as >99% of reconstructed sentences were identical to the original sentences. The text-embedding-ada-002 ada model uses a text embedding size of 1536, which is substantially larger than our image embedding size of 16 and also larger than the number of characters in the text phenotypes. Consequently, in this experiment, the text latent space encoding step does not represent dimensionality reduction, but rather encoding text into a latent space which is capable of representing a much larger range of text than the experimental text phenotypes. In the genome-to-embedding prediction step, text embeddings, similar to image embeddings, showed moderate heritability and prediction accuracy on the test set, with an average accuracy of 0.27 and a standard deviation of 0.05.

#### Comparing genome-to-text prediction and conventional approaches

Genome-to-text prediction from SNP markers produced text with a realistic vocabulary. End-to-end prediction accuracy of predicted text was measured as squared Pearson correlation *r*^2^ between true and predicted values for two extracted traits. End-to-end prediction accuracy for genome-to-text prediction was nonzero, although it did not reach conventional genomic prediction accuracy for either trait (see Table 3).

**Table 3.** Comparison between accuracy (r^2^) for end-to-end genome-to-text prediction and conventional genomic prediction for extracted traits.

|  | Prediction accuracy (sd) |  |
| --- | --- | --- |
|  | end-to-end | conventional |
| Length | 0.086 (0.024) | 0.188 (0.083) |
| Redness | 0.220 (0.025) | 0.329 (0.074) |

Figure 5 shows the difference between end-to-end prediction accuracy (r^2^) and conventional genomic prediction accuracy (r^2^) for genome-to-text prediction for two extracted traits. Each point within a distribution represents the difference calculated for a single experimental run (out of 50) on one trait. A difference of less than 0 indicates that end-to-end genomic prediction performed worse than conventional genomic prediction for that trait on that run, while difference greater than 0 indicates that end-to-end genomic prediction performed better than conventional genomic prediction for that trait on that run. End-to-end prediction accuracy was slightly lower numerically than conventional genomic prediction accuracy, but not significantly so.

**Figure 5.**
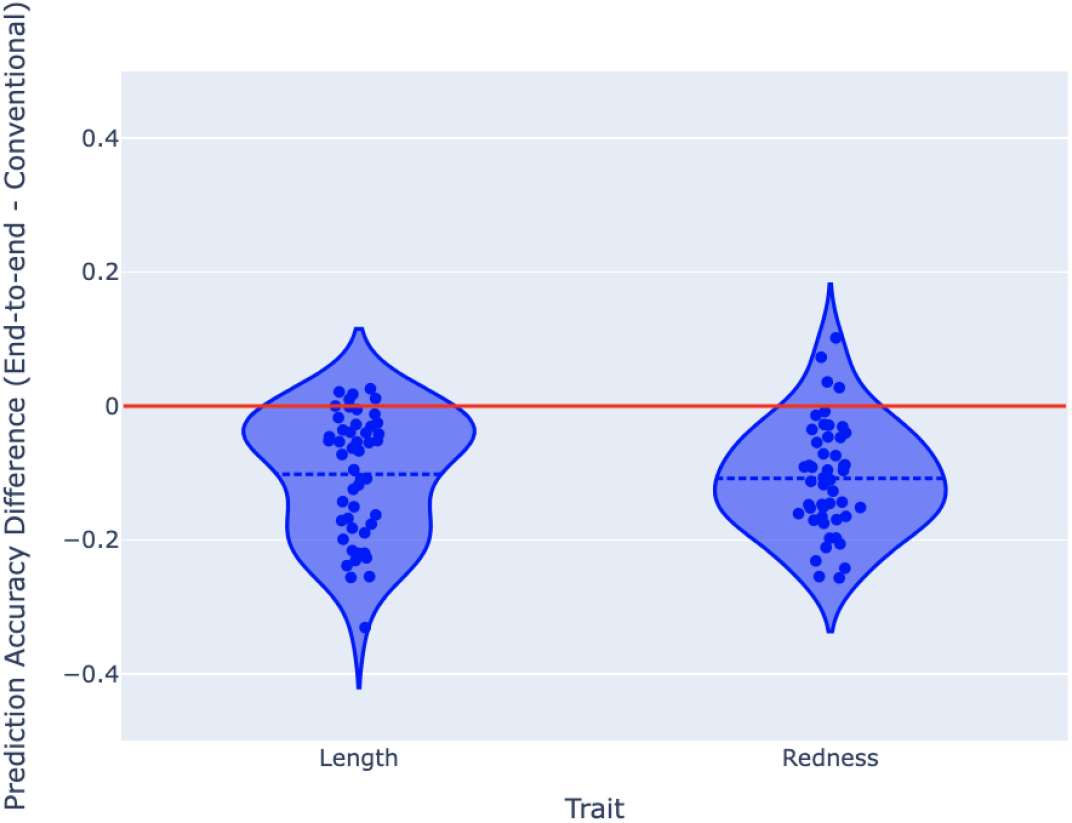
Violin plot of the difference between end-to-end prediction accuracy (r^2^) and conventional genomic prediction accuracy (r^2^) for genome-to-text prediction, as measured by two extracted traits.

#### Interpreting the genome-to-text prediction pipeline across genotypes

Table 4 compares the outputs from each step of our genome-to-text method. As with Figure 4 for genome-to-image, columns represent four different stages in our end-to-end method: here they represent original text, decoded mean embedding text, mean decoded mean embedding text, and end-to-end predicted text. The rows represent one clone each and comprise three of the four clones selected for Figure 4 (the lower resolution in text phenotypes compared to image phenotypes meant that a fourth genotype could not be distinguishable from all others on the average phenotype level). Each sentence in the “Original” column represents one strawberry fruit and corresponds to an image in Figure 4, in order. Compared to the image phenotypes, some information is lost, as the full range of variability is not able to be captured by the limited vocabulary for shape (“long”, “short”, “medium”) and color (“deep red”, “light red”), and most text descriptions do not contain information on fruit tapering. However, differences between clones are still apparent from the text phenotypes; for example, the first clone is generally longer and redder than the second clone, which is reflected by a higher use of the word “short” and a higher use of “light red” in the second clone compared to the first clone. The third clone is lighter than the first clone but longer than the second clone. The “decoded mean embedding” column shows the results of encoding each original sentence using text-embedding-ada-002 and decoding the embedding using vec2text. Reconstruction accuracy was extremely high, as the decoded mean embedding text descriptions as shown by the near perfect resemblance between original and decoded mean embedding descriptions. Text embeddings were averaged to produce one mean embedding for each clone, which can be decoded to the text in the “Mean decoded mean embedding” column. In the “End-to-End” column, we show the decoded text resulting from the SNP-predicted embedding for each clone. For the first and third clones, the predicted text corresponds perfectly to the true text, while for the second clone, the correct word for shape (“short”) was predicted, but a nonsensical combination of “long” and “short” was combined in the predicted sentence “Long, light red short strawberry.”. This moderate to low, but nonzero, prediction accuracy subjectively visible in Table 4 is consistent with the prediction accuracy of 0.086 for length and 0.222 for redness that we observed (Table 3).

**Table 4.** Comparison of original, decoded mean embedding, and end-to-end strawberry text descriptions.

| Clone | Fruit | Original | Decoded embeddings | Decoded mean embedding | End-to-end |
| --- | --- | --- | --- | --- | --- |
| 1 | 1 | Long, deep red strawberry. | Long, deep red strawberry. | Long, deep red strawberry. | Long, deep red strawberry. |
|  | 2 | Long, deep red strawberry. | Long, deep red strawberry. |  |  |
|  | 3 | Short, deep red strawberry. | Short, deep red strawberry. |  |  |
|  | 4 | Long, light red strawberry. | Long, light red strawberry. |  |  |
|  | 5 | Long, deep red strawberry. | Long, deep red strawberry. |  |  |
|  | 6 | Long light red strawberry. | Long light red strawberry. |  |  |
|  | 7 | Long, deep red strawberry. | Long, deep red strawberry. |  |  |
|  | 8 | Long, light red strawberry. | Long, light red strawberry. |  |  |
|  | 9 | Long, deep red strawberry. | Long, deep red strawberry. |  |  |
|  | 10 | Short, deep red strawberry. | Short, deep red strawberry. |  |  |
|  | 11 | Long, deep red strawberry. | Long, deep red strawberry. |  |  |
|  | 12 | Long, light red strawberry. | Long, light red strawberry. |  |  |
|  | 13 | Long, light red strawberry. | Long, light red strawberry. |  |  |
|  | 14 | Long, deep red strawberry. | Long, deep red strawberry. |  |  |
|  | 15 | Short, deep red strawberry. | Short, deep red strawberry. |  |  |
|  | 16 | Short, light red strawberry. | Short, light red strawberry. |  |  |
|  | 17 | Long and deep red. | Long and deep red. |  |  |
|  | 18 | Long, deep red strawberry. | Long, deep red strawberry. |  |  |
|  | 19 | Deep red, short strawberry. | Deep red, short strawberry. |  |  |
|  | 20 | Deep red, elongated shape. | Deep red, elongated shape. |  |  |
| 2 | 1 | Long, light red strawberry. | Long, light red strawberry. | Short, light red strawberry. | Long, light red short strawberry. |
|  | 2 | Short, pale strawberry. | Short, pale strawberry. |  |  |
|  | 3 | Medium length, light red. | Medium length, light red. |  |  |
|  | 4 | Short, light red strawberry. | Short, light red strawberry. |  |  |
|  | 5 | Short, light red strawberry. | Short, light red strawberry. |  |  |
|  | 6 | Short, pale red strawberry. | Short, pale red strawberry. |  |  |
|  | 7 | Short, light red strawberry. | Short, light red strawberry. |  |  |
|  | 8 | Long, light red strawberry. | Long, light red strawberry. |  |  |
|  | 9 | Long, light red strawberry. | Long, light red strawberry. |  |  |
|  | 10 | Pale, short strawberry. | Pale, short strawberry. |  |  |
|  | 11 | Short, light red, bluish. | Short, light red, bluish. |  |  |
|  | 12 | Short, light red strawberry. | Short, light red strawberry. |  |  |
|  | 13 | Pale, medium-length strawberry. | Pale, medium-length strawberry. |  |  |
|  | 14 | Long, light red strawberry. | Long, light red strawberry. |  |  |
|  | 15 | Short, pale red strawberry. | Short, pale red strawberry. |  |  |
|  | 16 | Long light red strawberry. | Long light red strawberry. |  |  |
|  | 17 | Medium long, light red. | Medium long, light red. |  |  |
|  | 18 | Medium-sized, light red strawberry. | Medium-sized, light red strawberry. |  |  |
|  | 19 | Long light red strawberry. | Long light red strawberry. |  |  |
|  | 20 | Long, light red strawberry. | Long, light red strawberry. |  |  |
| 3 | 1 | Long, light red strawberry. | Long, light red strawberry. | Long, light red strawberry. | Long, light red strawberry. |
|  | 2 | Long, light red strawberry. | Long, light red strawberry. |  |  |
|  | 3 | Long, light red strawberry. | Long, light red strawberry. |  |  |
|  | 4 | Long, light red strawberry. | Long, light red strawberry. |  |  |
|  | 5 | Long light red strawberry. | Long light red strawberry. |  |  |
|  | 6 | Long, light red strawberry. | Long, light red strawberry. |  |  |
|  | 7 | Deep red, long strawberry. | Deep red, long strawberry. |  |  |
|  | 8 | Long, deep red strawberry. | Long, deep red strawberry. |  |  |
|  | 9 | Long, light red strawberry. | Long, light red strawberry. |  |  |
|  | 10 | Long, light red strawberry. | Long, light red strawberry. |  |  |
|  | 11 | Long, light red strawberry. | Long, light red strawberry. |  |  |
|  | 12 | Long, light red strawberry. | Long, light red strawberry. |  |  |
|  | 13 | Short, deep red strawberry. | Short, deep red strawberry. |  |  |
|  | 14 | Deep red, elongated fruit. | Deep red, elongated fruit. |  |  |
|  | 15 | Long, deep red strawberry. | Long, deep red strawberry. |  |  |
|  | 16 | Long, deep red strawberry. | Long, deep red strawberry. |  |  |
|  | 17 | Long, deep red strawberry. | Long, deep red strawberry. |  |  |
|  | 18 | Long, light red strawberry. | Long, light red strawberry. |  |  |
|  | 19 | Long, deep red strawberry. | Long, deep red strawberry. |  |  |
|  | 20 | Long, deep red strawberry. | Long, deep red strawberry. |  |  |

#### Benchmark

Our validation also showed promising computational efficiency, with the entire process requiring under 2 hours per replicate on a T4 GPU with 51 GB of RAM.

## Discussion

In this study, we have introduced a new approach for predicting images and text from genetic markers, which we have named “end-to-end genomic prediction”. This approach combines nonlinear latent space encoding of high-dimensional phenotypes such as images and text using neural networks with linear prediction of breeding values. In our proof-of-concept experiments using a strawberry dataset, we show for the first time that genomic prediction of images and text can reach prediction accuracy comparable to that of conventional genomic prediction, using scalar numeric traits extracted from images and text as a mode of comparison. Nonetheless, there remains room for improvement in these methods.

For both images and text, we believe the latent space encoding step can be improved. For genome-to-image prediction, the latent space encoding step relied on a variational autoencoder that we trained on a subset of the strawberry fruit images in our dataset. The trained autoencoder showed only moderate reconstruction accuracy for shape and color traits, including notable blurring of small-scale features. Identifying neural network architectures that preserve a higher level of detail with greater accuracy is one area for improvement, but it will also be important to verify that detail encoded in embeddings is also heritable. For genome-to-text prediction, reconstruction accuracy was very high, but this did not translate to high end-to-end accuracy for shape and color. Reconstruction accuracy and embedding prediction accuracy jointly determine the accuracy of end-to-end genomic prediction, but improving these two factors alone may not be sufficient to ensure high end-to-end prediction accuracy. One desirable property of latent space embeddings which could contribute to end-to-end accuracy outside of reconstruction accuracy is linear interpolability, which is not a guaranteed property of the variational autoencoder approach we used^[2,35]^. Additionally, adding more embedding components does not necessarily improve prediction performance, particularly when those embedding components have low heritability. More work needs to be done to systematically identify the properties that image or text embeddings need to work effectively with genomic prediction.

While substantial variability in shape and color exists in the image dataset we used, the variability within each clone is relatively high compared to the variability between clones. Figure 6 shows the outputs for each stage of end-to-end genomic prediction for four test set clones which were not picked for high variability, unlike in Figure 4. While the differences in end-to-end predictions (“End-to-end” column) are rather small, it is worth noting that this can be explained by the low variability in the true averaged image phenotypes (“Mean encoded-decoded” column). Correspondingly, the variability in the conventional genomic predictions of extracted traits and their true values are both relatively low. This indicates that while low heritability affects the accuracy of end-to-end genomic predictions, it is no more of a constraint than it is with conventional genomic prediction methods. Some arbitrary differences in rotation exist between images, including a few cases of images that are upside down according to our calyx-up convention. As image rotation is not heritable and has a substantial effect on the image matrix, these cases may have a negative effect on end-to-end prediction accuracy. Either improving our preprocessing code to improve standardization of image rotation or using an autoencoder that can accommodate different image rotations could lead to improved end-to-end prediction accuracy. Additionally, while we predicted breeding values of images and text in our experiment, other applications of end-to-end genomic prediction may require predictions of total genetic value of image or text phenotypes.

**Figure 6.**
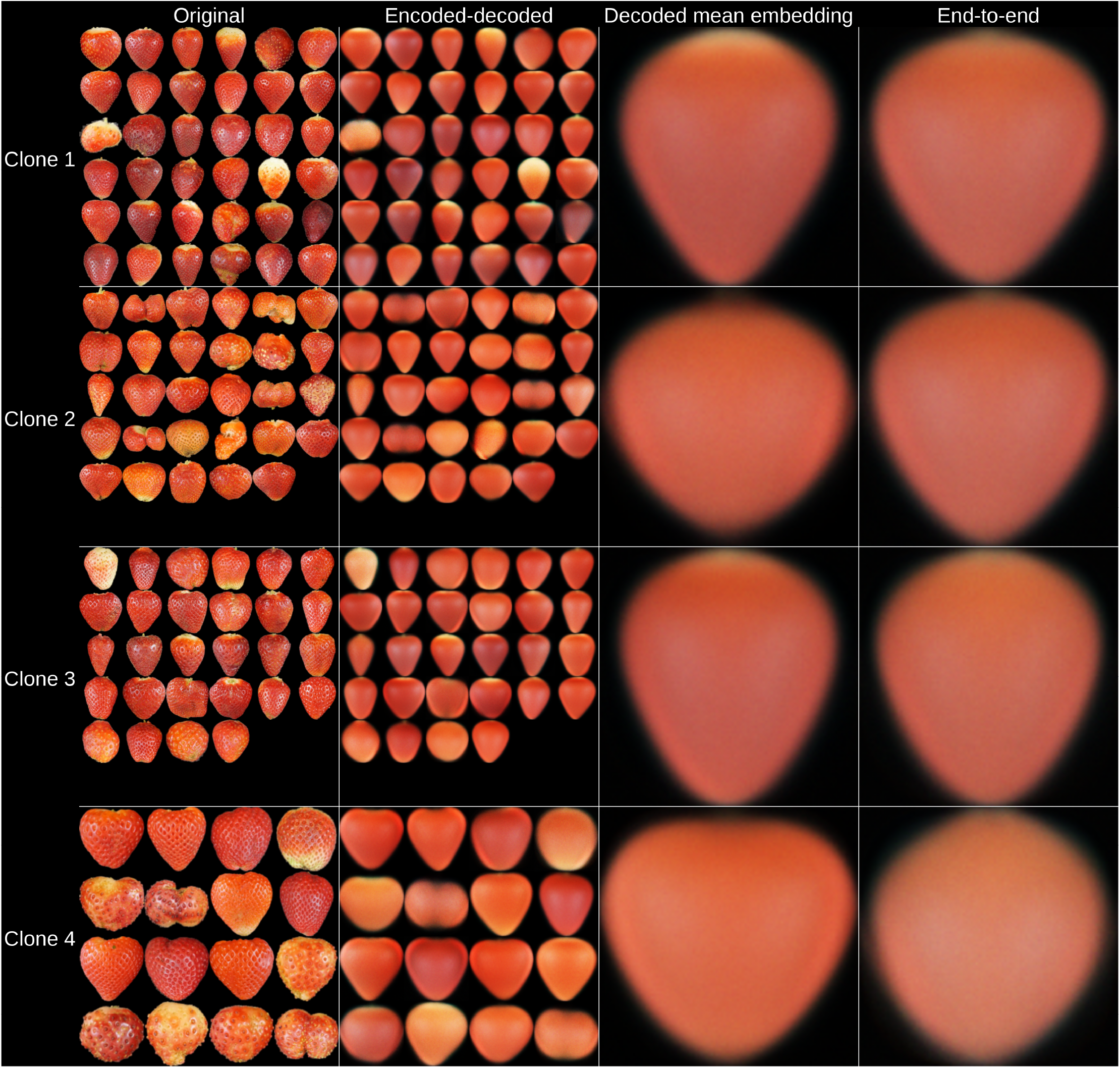
A comparison between genome-to-image outputs for four clones in the test set. The four columns represent original images, encoded-decoded images, mean encoded-decoded image, and end-to-end predicted image. Each row represents one strawberry clone.

While we used scalar numeric phenotypes extracted from image and text phenotypes as a means to compare end-to-end genomic prediction with conventional genomic prediction, it is not necessary to know what features contribute to the differences between images in order to perform accurate end-to-end genomic prediction of images or text. In our method, neither the latent space encoding step nor the embedding prediction step requires extracted traits or information specifying which extracted traits are relevant. Because our method does not require preexisting knowledge about extracted traits, we believe it has the potential to be generalizable across a wider range of plant traits and species than we tested, for example in fruit and foliage aesthetic selection, in visual disease scoring, or in food crop sensory evaluations. End-to-end genomic prediction could also benefit animal breeding, where traits such as body conformation, structural soundness, locomotion, and even behavioral descriptions are more naturally represented as images or text. Applying end-to-end genomic prediction in other crops and in livestock could therefore create opportunities to improve selection for traits that have traditionally relied on subjective scoring or costly measurements. Additionally, in contrast with previous methods for image prediction in breeding^[22]^, our method effectively uses genome-wide markers, without the need for pre-filtering markers based on association with extracted image or text features, enabling a truly “end-to-end” approach. This simplifies the tasks of genome-to-image prediction and genome-to-text prediction to require only genotypes and their respective image or text phenotypes for training. Consequently, we see potential for end-to-end genomic prediction in breeding where selections are made based on “the breeder’s eye”: complex aesthetic or sensory traits that have not been or cannot be successfully decomposed into a small set of numeric features. In these cases, prediction of images or text could make the improved selection accuracy and generation length which genomic selection enables, without simplifying phenotypes to potentially insufficient components.

## Availability of Source Code

Python and R code to reproduce these analyses are available on GitHub^[39]^. Code is structured as Jupyter notebooks designed to be run in a Google Colab T4 runtime, as explained in the provided README file.

## Availability of Supporting Data

The data supporting the results of this article are available in Zenodo^[40]^.

## Abbreviations

GP: genomic prediction
LWR: length-width ratio
MSE: mean squared error
SNP: single-nucleotide polymorphism
VAE: variational autoencoderr

## ACKNOWLEDGEMENTS

We thank Dr. Gota Morota for his comments and advice on the work described in this manuscript.

